# The Role of Nde1 Phosphorylation in Interkinetic Nuclear Migration and Schizophrenia

**DOI:** 10.1101/2024.02.25.581986

**Authors:** David J Doobin, Paige Helmer, Aurelie carabalona, Chiara Bertipaglia, Richard B Vallee

## Abstract

Nde1 is a cytoplasmic dynein regulatory protein with important roles in vertebrate brain development. One noteworthy function is in the nuclear oscillatory behavior in neural progenitor cells, the control and mechanism of which remain poorly understood. Nde1 contains multiple phosphorylation sites for the cell cycle-dependent protein kinase CDK1, though the function of these sites is not well understood. To test their role in brain development we expressed phosphorylation-state mutant forms of Nde1 in embryonic rat brains using *in utero* electroporation. We find that Nde1 T215 and T243 phosphomutants block apical interkinetic nuclear migration (INM) and, consequently, mitosis in radial glial progenitor cells. Another Nde1 phosphomutant at T246 also interfered with mitotic entry without affecting INM, suggesting a more direct role for Nde1 T246 in mitotic regulation. We also found that the Nde1 S214F mutation, which is associated with schizophrenia, inhibits CDK5 phosphorylation at an adjacent residue which causes alterations in neuronal lamination. These results together identify important new roles for Nde1 phosphorylation in neocortical development and disease and represent the first evidence for Nde1 phosphorylation roles in INM, neuronal lamination, and schizophrenia.

**Significance Statement:** The results presented in this study show the importance of Nde1 phosphoregulation during successive stages of neocortical development. Nde1 dysregulation may, in turn, have important consequences in human psychiatric disorders, such as schizophrenia, as well. We observed clear and potent effects of specific Nde1 phosphomutants in the control of INM and neural development. Our results provide strong support for new insight into the little-understood mechanism for triggering RGP mitotic entry. In the course of these studies, we obtained clear evidence for a novel post-mitotic role for Nde1 phosphorylation in schizophrenia.

## Introduction

Neocortical development is a complex process involving multiple steps in neurogenesis and subsequent post-mitotic neuronal migration. In earlier work, we found that the minus end-directed microtubule motor protein cytoplasmic dynein, and a number of its regulatory proteins, play prominent roles in these behaviors.^1–3^ The dynein regulators include LIS1, its interactors Nde1 and Ndel1, and the nuclear pore-binding proteins BicD2 and CENP-F, which are known to be involved in dynein recruitment to the nuclear envelope (NE) (Figure 1). Interference with dynein expression or function, or that of its regulatory proteins, causes a number of brain developmental defects. LIS1 haploinsufficiency causes type I lissencephaly,^3–5^ mutations in Nde1 cause severe forms of microcephaly or microlissencephaly, and mutations in Ndel1 are embryonic lethal.^3,6–9^

**Figure 1.**
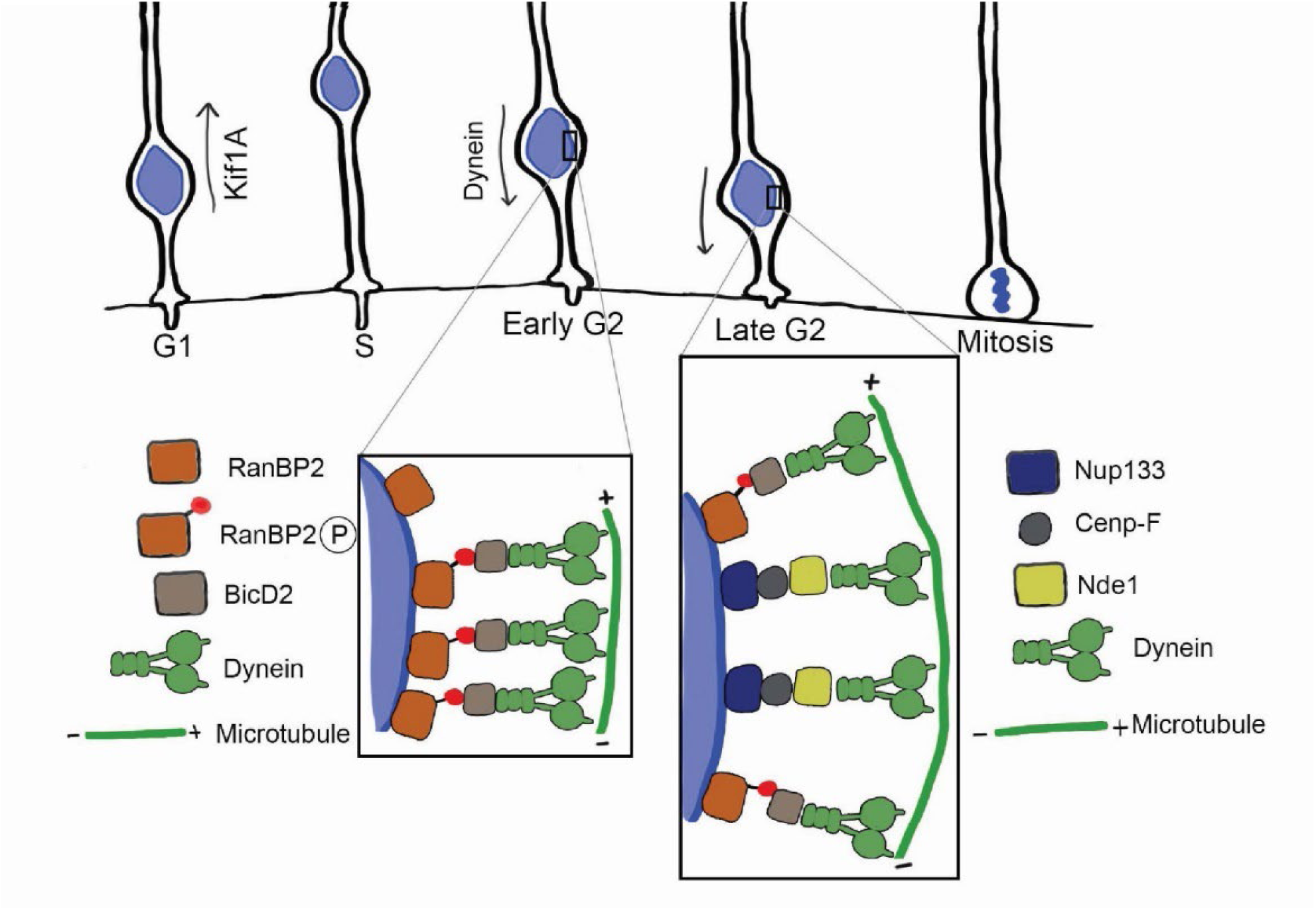
Schematic Representation of Nde1 roles in Neocortical Development. Our previous work found Nde1 to be involved in two aspects of INM: G2-dependent apical nuclear migration and mitotic entry at the ventricular surface of the brain. Apical INM is under the control of the kinesin Kif1A and basal INM is under Dynein control. Dynein is recruited to the nuclear envelope (NE) in two stages: in early G2, phosphorylated RanBP2 bind BicD2, which binds Dynein to the NE, while in late G2, a scaffold of Nup133-CENP-F-Nde1 recruits Dynein. Once the nucleus reaches the ventricular surface, mitosis occurs. The mechanisms by which Nde1 functions in both INM and in mitotic entry are unknown.

Previous work in our lab revealed that reduced Nde1 expression causes a marked decrease in mitotic index in the radial glial progenitor cells (RGPs),^3^ which give rise to most of the neurons in the developing neocortex.^10^ The RGP somata are located within the ventricular zone (VZ) of the developing brain, but also extend highly elongated basal processes spanning the distance from the ventricular to the pial surfaces of the brain and providing scaffolding for neuronal migration and cortical assembly.^10^ RGP nuclei undergo a remarkable form of cell cycle-dependent oscillatory behavior known as interkinetic nuclear migration (INM). During G1, the nucleus travels basally, away from the ventricle, and then, during G2, apically back to the ventricle (Figure 1). We found these behaviors to be driven, respectively, by the kinesin Kif1A, and cytoplasmic dynein.^1^ We also found that, for mitosis to proceed, the RGP nucleus must reach the apical terminus of the RGP cell, where the centrosome resides.^11^ Whether the juxtaposition of centrosome and nucleus is sufficient for mitotic entry or if other conditions must be fulfilled remains uncertain.

Apical INM in RGPs is activated by the G2-specific protein kinase CDK1, which controls dynein recruitment to the RGP nuclear envelope.^2,12^ One role for CDK1 is to phosphorylate the nucleoporin RanBP2 during early G2,^13^ which activates its recruitment of the dynein adaptor BicD2, and, in turn, dynein itself, to the NE.^11,13,14^ As G2 progresses, Cdk1 also phosphorylates Nde1 to activate its recruitment of cytoplasmic dynein to the large NE- and kinetochore-binding scaffold protein CENP-F (Figure 1).^11,15^ This behavior occurs when CENP-F emerges from inside the nucleus and binds to the NE via the nucleoporin Nup133.^16^

These observations and studies in non-neuronal cells^13–15^ have suggested that BicD2-mediated dynein recruitment temporally precedes CENP-F-mediated dynein recruitment, presumably resulting in a gradual increase in apically directed dynein-generated force at with the RGP NE as G2 progresses.^11^ Consistent with this hypothesis, BicD2 expression can rescue INM in both BicD2- or CENP-F knockdown conditions.^11^ Curiously, however, BidD2 could not rescue a defect in RGP mitotic entry caused by Nde1 knockdown.^3^ This result suggested that Nde1 might play a unique additional role directly in RGP mitotic entry.

In contrast to the clear effects of Nde1 RNAi we have observed on RGP behavior,^3^ we found little effect in these cells of RNAi for the Nde1 paralogue Ndel1, but instead a clear inhibitory effect on post-mitotic neuronal migration.^3^ The differences in cellular behavior associated with the two orthologues presumably account for observed phenotypic differences *in vivo*. Human NDE1 mutations cause decreased neurogenesis,^3,6,8^ manifesting in a hereditary form of microcephaly, with little reported effect on post-mitotic neuronal behavior. In contrast, a mouse model with a Ndel1 deletion exhibited microlissencephaly,^17^ consistent with a more substantial defect in post-mitotic neuronal migration.

Nde1 and Ndel1 each contain multiple sites for CDK1 and CDK5 phosphorylation, though the relative contributions of these sites in controlling cellular and subcellular behavior are incompletely known in general, and in the developing brain in particular. Mutations in Ndel1 at its Cdk1 and Cdk5 phosphorylation sites have been reported to reduce Ndel binding to the G2 NE.^18^ Effects of Ndel1 phosphorylation on lysosome transport have also been reported.^19,20^ We more recently found pronounced effects of Cdk1 phosphorylation of Nde1 on nucleus and kinetochore recruitment in dividing Hela cells.^15^ Cdk1 phosphorylation enhanced Nde1 recruitment to the prophase NE and markedly prolonged Nde1 localization at kinetochores, surprisingly into late anaphase. This pattern was strikingly similar to that of the Nde1 interactor CENP-F. Overexpression of wild type (WT) NDE1 or of a triple CDK1 phosphomimetic mutant also resulted in clear decoration of the G2 NE in HeLa cells, while triple phosphomutant NDE1 failed to localize to the NE.^15^

Previous studies focused on three residues as phosphorylation sites within the disordered region of Nde1: T215, T243, and T246, all predicted CDK1 phosphorylation sites. Other potential CDK1 sites in Nde1 are poorly conserved between species, and deemed less likely to be of physiological interest for the current study. Given the previous work establishing these three sites in the recruitment of NDE1 to the NE and the poor conservation of other CDK1 sites, this study focuses on T215, T243, and T246 as potential regulators of NDE1 function in RGPs.

In addition to a role in microcephaly, mutations in NDE1 have been associated with a spectrum of neuropsychiatric diseases, including schizophrenia.^21–23^ Several families with severe microcephaly or microlissencephaly have been found to have homozygous recessive deletions of NDE1 that result in either premature truncation of the protein, or no detectable expression.^8,9^ Copy number variants of NDE1 have also been linked to schizophrenia.^23^ Nde1 is also a known binding partner of DISC1, mutations in which were found to be responsible for the first known heritable form of schizophrenia.^21,24,25^ A more recent study has also identified six unrelated schizophrenia patients with a shared NDE1 missense mutation (S214F).^22^ Intriguingly, this mutation is adjacent to the NDE1 T215 residue, which is a known target of Cdk1 and Cdk5.^8,26^ How defects in NDE1 expression and phosphorylation might contribute to this disease is unknown.

The current study was initiated to determine the functions of Nde1 phosphorylation in the developing brain. We present the first data demonstrating a role for phosphorylation at each of three majors predicted Nde1 CDK1 sites in the developing rat neocortex, T215, T243, and T246. Importantly, these data provide clear support for a distinct, direct role for Nde1 phosphorylation in controlling the classic RGP nuclear oscillations comprising interkinetic nuclear migration. In addition, we identify a single Nde1 phosphorylation site, Nde1 T246, which directly controls mitotic entry. Finally, we provide evidence that Nde1 phosphorylation at a predicted CDK1 residue is controlled by mutation of the adjacent schizophrenia-associated Nde1 residue, a novel experimental link between a cytoplasmic dynein regulatory gene and a neuropsychiatric disorder.

## Results

To determine the physiological effects of Nde1 phosphorylation in brain development, we first focused on the three Cdk1 sites: T215, T243, and T246. Of these sites, T243 is also a predicted Cdk5 target. We performed *in utero* electroporation in embryonic day 16 (E16) rat embryos by injecting either GFP or Nde1 shRNA in combination with mCherry-tagged Nde1 constructs into the left cortical ventricle. Expression of GFP or of Nde1 shRNA alone were used as controls. At E20 or postnatal day 6 (P6), brains were fixed and examined for effects of these constructs on neocortical development. The apical process length in the RGPs was determined by measuring the distance from the apical side of the nucleus to the ventricular surface. As consistent with previous work, Nde1 shRNA resulted in accumulation of nuclei between 10 and 15µm from the ventricular surface, so the results were calculated as the percent of RGPs with a measured apical endfoot length (AEL) of 15 µm or less. Mitotic index was calculated as the percentage of transfected cells that were phospho-Histone H3 (PH3) positive compared to the total number of transfected cells.

As previously observed,^3^ Nde1 RNAi caused accumulation of the majority of RGP nuclei between 10 and 15μm from the ventricular surface of the developing rat brain (Figure 2A-B). Co-expression of wild-type NDE1 engineered to resist RNA-inhibited expression restored the RGP nuclei to their expected distribution relative to the VS (Figure 2A-B). The mitotic index for the Nde1 shRNA-expressing RGP cells was also markedly reduced, an effect that could be rescued by co-expression of wild-type NDE1 (Figure 2C), in agreement with the results of previously published studies.^3^ We also performed Nde1 RNAi rescue with a triple phosphomutant construct (T215A, T243A, and T246A, referred to here as Nde1x3A). As the fluorophore in this and other experiments shown (Figures 2,3,4,6) is directly conjugated to Nde1, the signal is visible as small puncta in Figure 2Aiii’-2Aiv’’’ The Nde1x3A construct failed to restore either INM or mitotic index, resulting in a distribution of RGP nuclei similar to that caused by Nde1 RNAi alone (Figure 2A-C). Together, these results suggest that INM, and as a consequence, RGP cell proliferation, are regulated by Nde1 phosphorylation.

**Figure 2.**
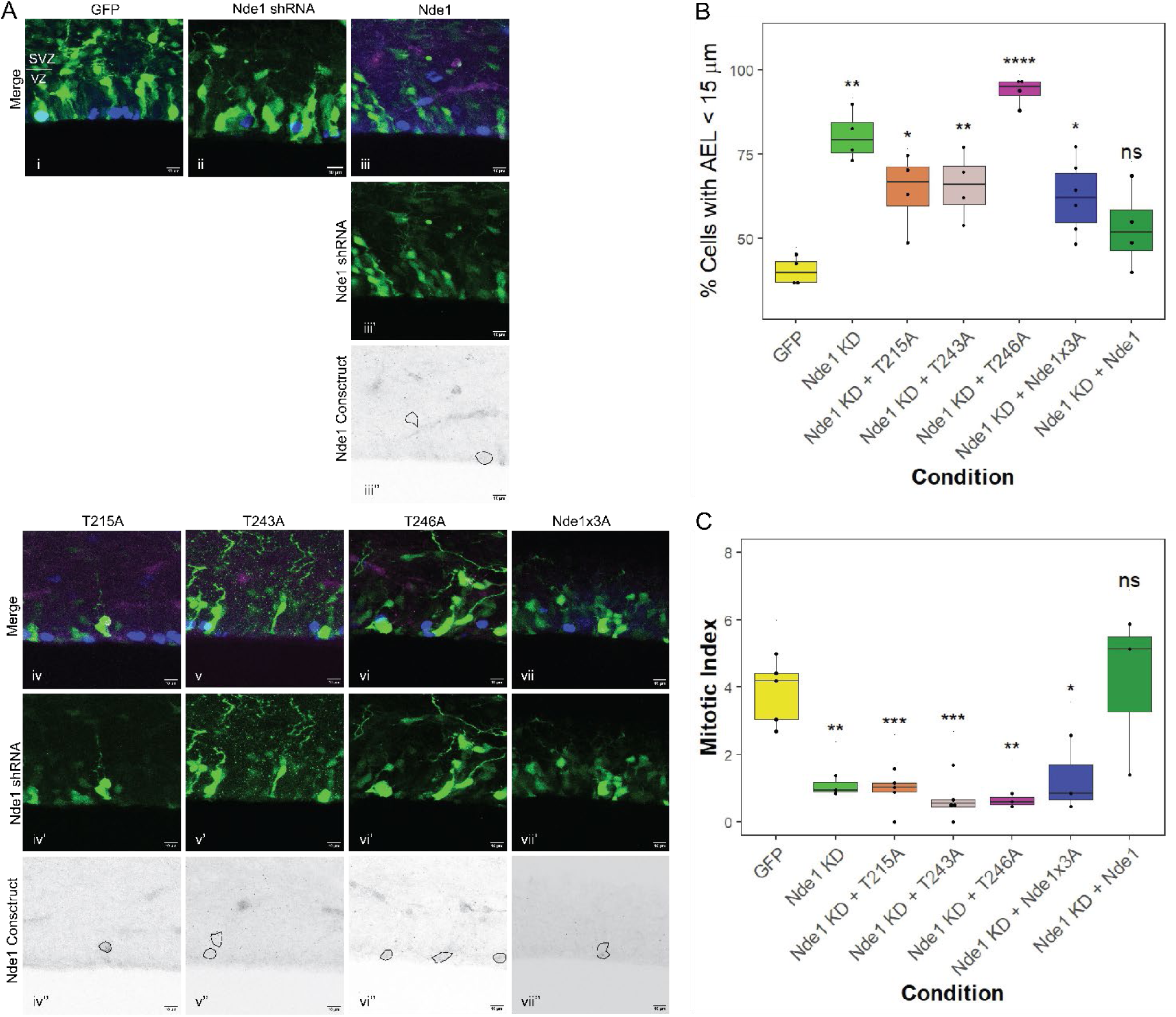
Rescue of Nde1 KD with Phosphomutant Constructs. 2A) Representative images of the VZ in brains injected with GFP (i), Nde1 KD (ii), Nde1 KD rescued with WT Nde1 (iii); and Nde1 KD rescued with single site phosphomutants T215A (iv), T243A (v), T246A (vi), and triple phosphomutant (vii). In merged images (2Ai-vii), PH3 is shown in blue, GFP or Nde1 shRNA in green, and Nde1 constructs in magenta. Nde1 shRNA is shown alone in 2Aiii’-vii’. Nde1 constructs are shown in black in 2Aiii’’-vii’’, with example cells outlined in black. Scale bars for all images are 10µm. A white line in the GFP control image shows the border between the SVZ (above) and VZ (below). 2B) Comparison of percent of cells with an apical endfoot length (AEL) less than 15 µm in RGPs expressing Nde1 shRNA and phosphomutant forms of Nde1. Apical endfoot length (AEL) was measured from the apical base of the nucleus directly to the ventricular surface in each case. 2C) Mitotic index values determined for RGP cells expressing phosphomutants of Nde1 co-transfected with Nde1 shRNA. Mitotic index was determined as the percent of transfected cells within the VZ that were positive for PH3. All statistical comparisons are made to control data. One-way ANOVA and post hoc Dunnet test were used in 2B-C, ns = not significant, *p<0.05, **p<0.01, ***p<0.001, ****p<0.0001. n = mean of values across at least 3 different embryos from at least 2 different operations per condition. All experiments were done with E16 operations and E20 analysis in developing rat brains. 2B-C data plotted as interquartile range with 5-95% whisker range.

**Figure 3.**
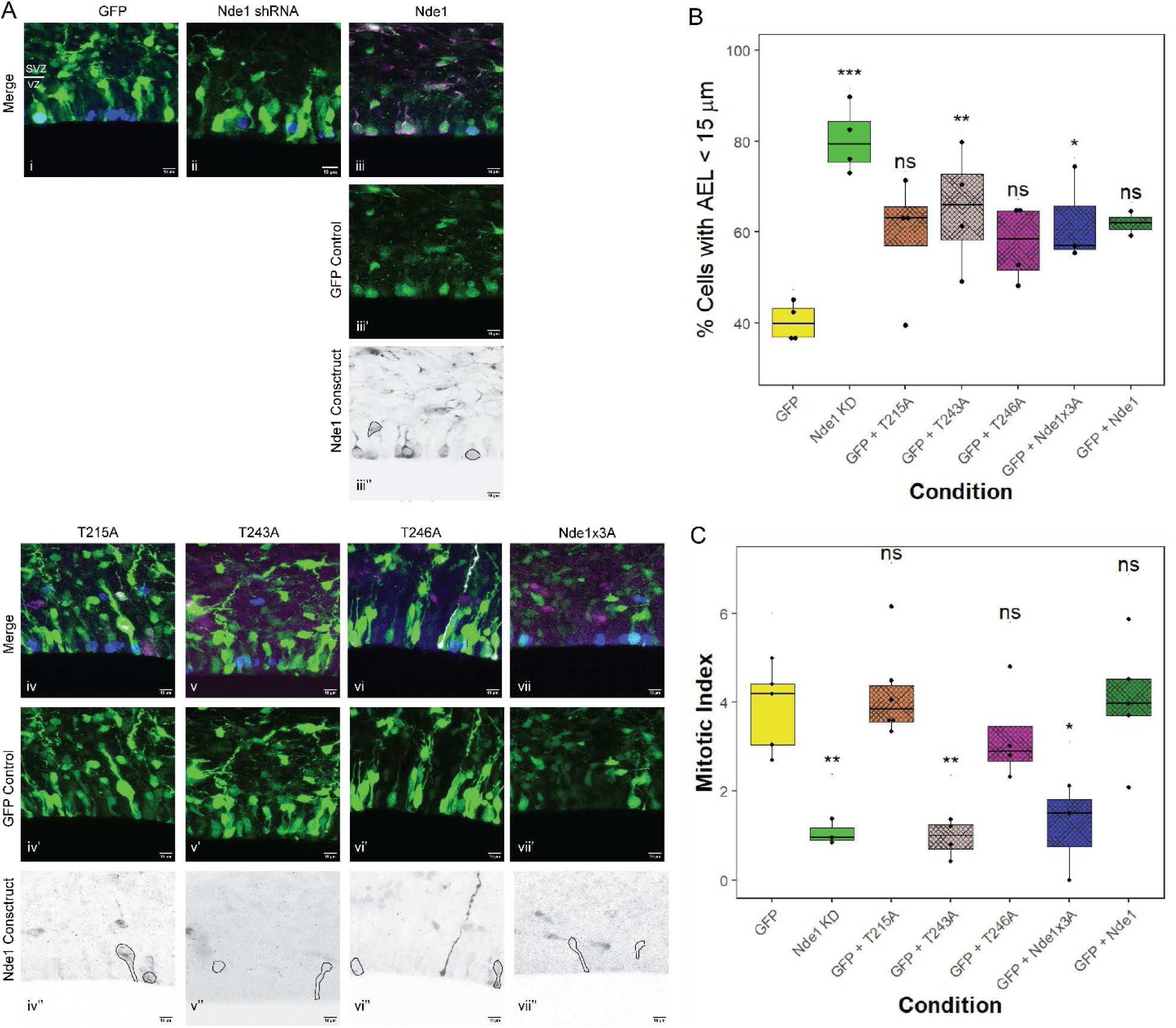
Dominant Negative Effects of Nde1 Phosphomutant cDNA. 3A) Representative images of RGP cells embryonic rat brain subjected to IUE with GFP control plasmid alone (i), Nde1 KD alone (ii), or GFP co-transfected with cDNAs expressing the Nde1 phosphomutants T215A (iii), T243A (iv), T246A (v) and Nde1x3A (vi). In merged images (3Ai-vii), PH3 is shown in blue, GFP or Nde1 shRNA in green, and Nde1 constructs in magenta. Nde1 shRNA is shown alone in 3Aiii’-vii’. Nde1 constructs are shown in black in 3Aiii’’-vii’’, with example cells outlined in black. Scale bars for all images are 10µm. A white line in the GFP control image shows the border between the SVZ (above) and VZ (below). 3B) Comparison of the percent of cells with an apical endfoot length (AEL) less than 15 µm in RGPs expressing GFP and phosphomutant forms of Nde1. 3C) Mitotic indices of phosphomutant constructs co-transfected with GFP. All statistical comparisons are made to control data. One-way ANOVA and post hoc Dunnet test were used in 3B-C), ns = not significant, *p<0.05, **p<0.01, ***p<0.001. n = mean of values across at least 3 different embryos from at least 2 different operations per condition. All experiments done with E16 operations and E20 analysis in developing rat brains. 3B-C) data plotted as interquartile range with 5-95% whisker range.

To determine the relative physiological importance of individual Nde1 phosphorylation sites, the effects of single-site phosphomutant constructs were also examined. For this purpose, we co-expressed a Nde1 shRNA along with RNAi-resistant versions of Nde1:T215A or T243A. Neither construct rescued the defect in INM caused by Nde1 shRNA. Instead, these treatments resulted in the accumulation of RGP nuclei between 10 and 15μm from the ventricular surface, similar to the effect seen of Nde1x3A expression or of Nde1 knockdown alone (Figure 2A-B). These conditions also resulted in a reduced mitotic index (Figure 2C). Interestingly, rescue of Nde1 RNAi with a mutation in the third site tested, T246A, resulted in accumulation of cells directly adjacent to the ventricular surface, though mitotic index was still reduced compared to the WT level (Figure 2A-C). These results suggest that Nde1-T246A can rescue INM, but not the G2-to-M transition, similar to the effects of Nde1 knockdown in combination with BicD2 rescue.^3^

We also tested for dominant negative effects of the Nde1 mutations (Figure 3). As Nde1 is dimeric, we expected that the phosphomutant form might interact with WT Nde1 molecules, resulting in a dimer that was less effective at recruiting dynein to the nucleus. This was tested by injecting an empty GFP reporter along with each Nde1 construct. In two cases - T243A, and Nde1x3A - we detected inhibition of INM (Figure 3A-B). Expression of T215A or T246A showed no effect on INM or mitotic index (Figure 3B-C). The only significant effect on mitotic index we observed was in T243A- and Nde1x3A-expressing cells (Figure 3C). In these cases, mitotic index was significantly reduced compared to control. As a control, we included WT Nde1 and GFP control, which showed no effect on INM or mitotic index (Figure 3Aiii, B-C)

In order to test the validity of the phosphorylation sites, we used a Nde1 construct that contained a mutation from T to glutamic acid, rather than alanine, in order to mimic the charge of a phosphorylated residue. We tested the effect of this construct (Nde1x3E) on INM and on mitotic index. We found that Nde1x3E had no effect on INM, either when co-transfected with Nde1 shRNA, or with GFP control (Figure 4B). This indicates that the phosphomimetic is able to rescue Nde1 function in INM, and that it has no dominant negative effect in the presence of endogenous Nde1. The mitotic index results, however, showed no effect in the GFP control and Nde1x3E condition, but did show a significant decrease in mitotic index when Nde1x3E was co-transfected with Nde1 shRNA (Figure 4C). This was an unexpected result, as all phosphomutants in combination with Nde1 shRNA resulted in a decrease in mitotic index. Only the T243A and Nde1x3A constructs had a dominant effect on mitotic index without Nde1 shRNA present. This may indicate that the three sites must be phosphorylated in a specific combination in order to trigger mitotic entry, and one or more sites must be dephosphorylated in order for this to occur. As T246A is able to rescue INM, but not mitotic index, it is likely that this residue is required to be phosphorylated in order for the cell to enter mitosis. T243A was the only construct that had an effect on INM and mitotic index even in the presence of endogenous Nde1, indicating that phosphorylation at this residue is required for INM. As INM is affected in this condition, it is possible that the mitotic index is affected as a result of RGP nuclei failing to reach the ventricular surface, so phosphorylation at T243 may be only indirectly required for mitosis to progress. Thus, either T215, T243, or both, may be required to be dephosphorylated in order to trigger mitotic entry at the ventricular surface. In order to fully interrogate this mechanism, a system would be required to selectively rescue INM without Nde1 function in order to specifically observe the effects of Nde1 phosphorylation on mitotic entry.

**Figure 4.**
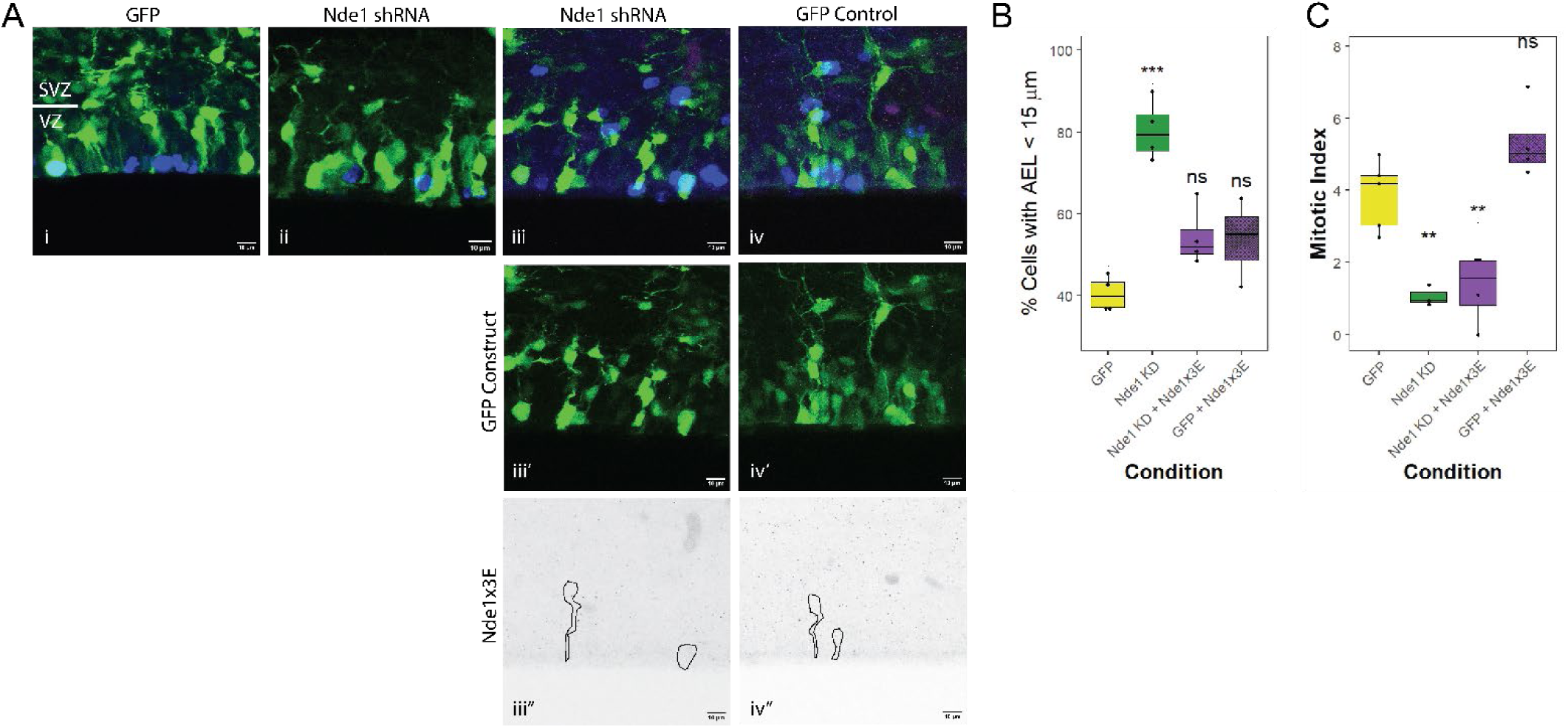
Effects of Nde1 phospho-mimetic constructs on INM and mitotic index. 4A) Representative images of RGP cells embryonic rat brain subjected to IUE with GFP control plasmid alone (i), Nde1 KD alone (ii), or Nde1x3E with Nde1 shRNA (iii) or GFP control (iv). In merged images (4Ai-iv), PH3 is shown in blue, GFP or Nde1 shRNA in green, and Nde1x3E in magenta. GFP or Nde1 shRNA is shown alone in 4Aiii’-iv’. Nde1x3E is shown in black in 3Aiii’’-iv’’, with example cells outlined in black. Scale bars for all images are 10µm. A white line in the GFP control image shows the border between the SVZ (above) and VZ (below). 4B) Comparison of the percent of cells an apical endfoot length (AEL) less than 15 µm of the ventricular surface in RGPs expressing GFP, Nde1 shNRA, or Nde1x3E with GFP or Nde1 shRNA. 4C) Mitotic indices of RGPs expressing GFP, Nde1 shNRA, or Nde1x3E with GFP or Nde1 shRNA. All statistical comparisons are made to control data. One-way ANOVA and post hoc Dunnet test were used in 4B-C), ns = not significant, *p<0.05, **p<0.01. n = mean of values across at least 3 different embryos from at least 2 different operations per condition. All experiments done with E16 operations and E20 analysis in developing rat brains. 4B-C) data plotted as interquartile range with 5-95% whisker range.

**Figure 5.**
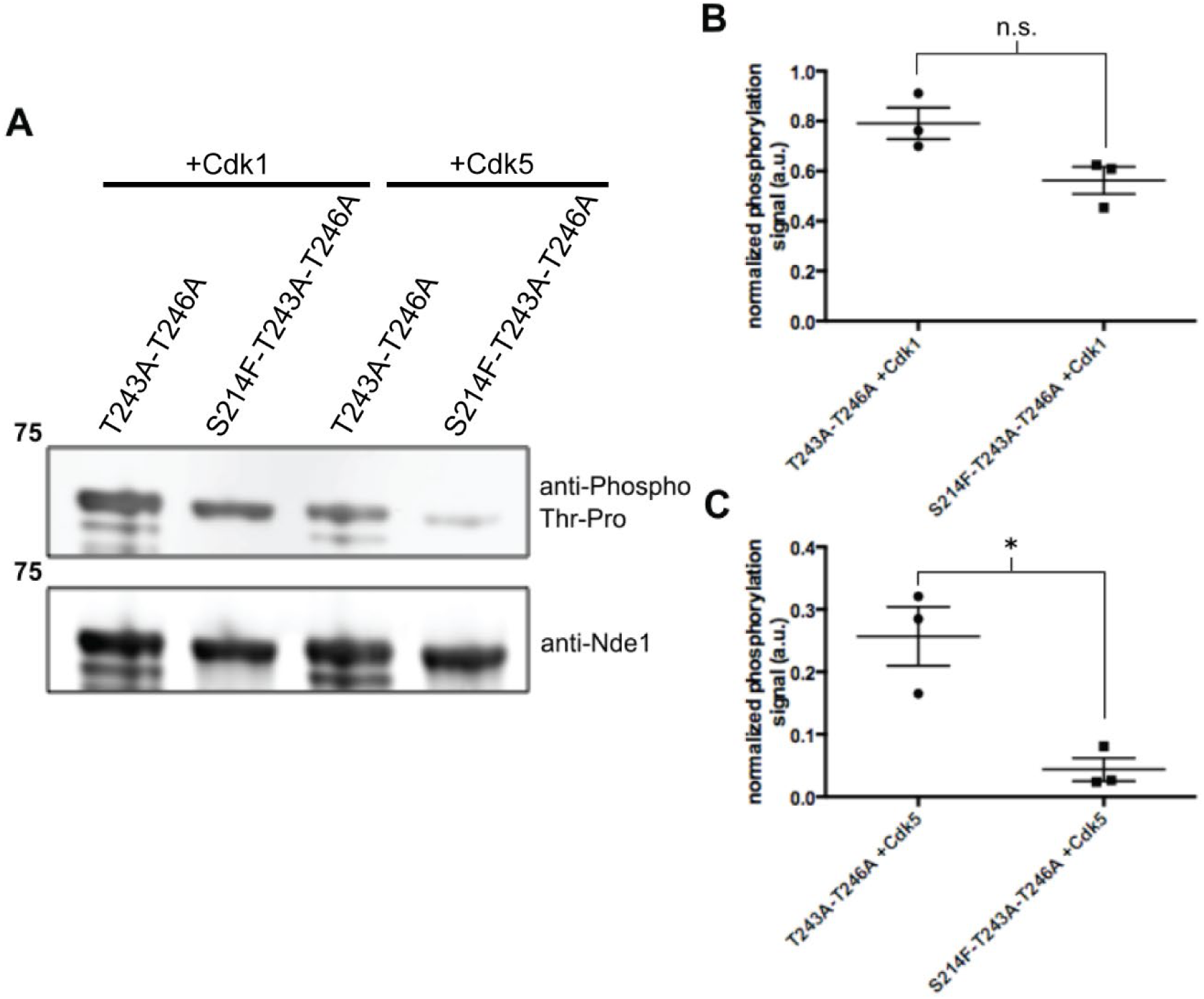
Phosphorylation of Nde1-S214F with Cdk1 and Cdk5. 5A) The human NDE1-encoding cDNA with the S214F mutation in tandem with phosphomutation of the T243 and T246 sites from T to A were expressed and purified to isolate the effect of the S214F mutation on Cdk1 and Cdk5 phosphorylation. Multiple bands are commonly seen for Nde1, a single band is likely seen in the S214F condition due to conformational changes in the protein due to the substitution of a large hydrophobic residue for an uncharged residue. There is a clear reduction in Cdk5 phosphorylation in Nde1 with the S214F mutation, but no effect on Cdk1 phosphorylation. 5B-C) Quantification of the anti-phosphothreonine signal relative to the anti-Nde1 signal. Data plotted as mean with standard deviation. Comparison using paired t-test, *p<0.01.

### Schizophrenia-associated Nde1 mutation

A missense mutation in NDE1, S214F, has been reported to be associated with a form of schizophrenia in a cohort of six unrelated Japanese patients.^22^ Nde1-S214 has been neither shown nor predicted to be phosphorylated. Nonetheless, to account for its particular pathophysiological effects, the S214F mutation was postulated to act in a highly unusual manner, sterically hindering phosphorylation of the adjacent T215 residue.^22^ To test this hypothesis we performed direct *in vitro* phosphorylation of recombinant NDE1 using commercially available CDK5.

We note that the S214F schizophrenia mutation replaces a polar hydrophilic amino acid with an aromatic, very hydrophobic one, which has a bulky side chain that may cause a marked effect on the local 3D structure of the polypeptide. We hypothesized that this conformational perturbation may affect Cdk5 accessibility to the adjacent T215 site, thereby reducing the ability of the protein kinase to phosphorylate Nde1. Because Cdk5 is also predicted to phosphorylate T243 and T246, we mutated these two residues from T to A before we expressed and purified GST-Nde1 full-length, with or without the S214F mutation (referred to as S214F-T243A-T246A and T243A-T246A). We then performed an *in vitro* kinase assay with recombinant Cdk5 and detected a 90% decrease in Nde1 phosphorylation associated with the NDE1 S214F schizophrenia mutation. Phosphorylation was assessed by quantitative immunoblotting of the recombinant Nde1-expressing cells using an anti-phospho-threonine antibody (Figure 4A, C). We found the presence of Nde1-S214F to cause a small, but non-significant, decrease in the level of Nde1 phosphorylation by Cdk1 (Figure 4A-B). These results suggest therefore, that mutation of the Nde1 S214 residue inhibits Cdk5, but not Cdk1, phosphorylation of T215.

To test the effects of the apparent specific inhibition of Cdk5 in S214F constructs, GFP was co-transfected with mCherry tagged Nde1-S214F in E16 rat brains (Figure 6). As all six known patients with the S214F mutation are heterozygous for this mutation, we did not perform concurrent knockdown of Nde1 with expression of the S214F mutant. Electroporation with Nde1-S214F cDNA had no effect on the distribution of RGP nuclei relative to the VS, or on RGP mitotic index by E20 (Figure 6B-C). As Cdk5 is preferentially expressed in post-mitotic neurons,^27^ and not in RGPs, this result is consistent with the *in vitro* kinase assay. If Cdk1 phosphorylation of Nde1 was affected by the S214F mutation, we might expect to see altered INM and mitotic entry in RGPs, but as Cdk5 phosphorylation appears to be specifically inhibited, we might expect to see results only in cells that express Cdk5.

**Figure 6.**
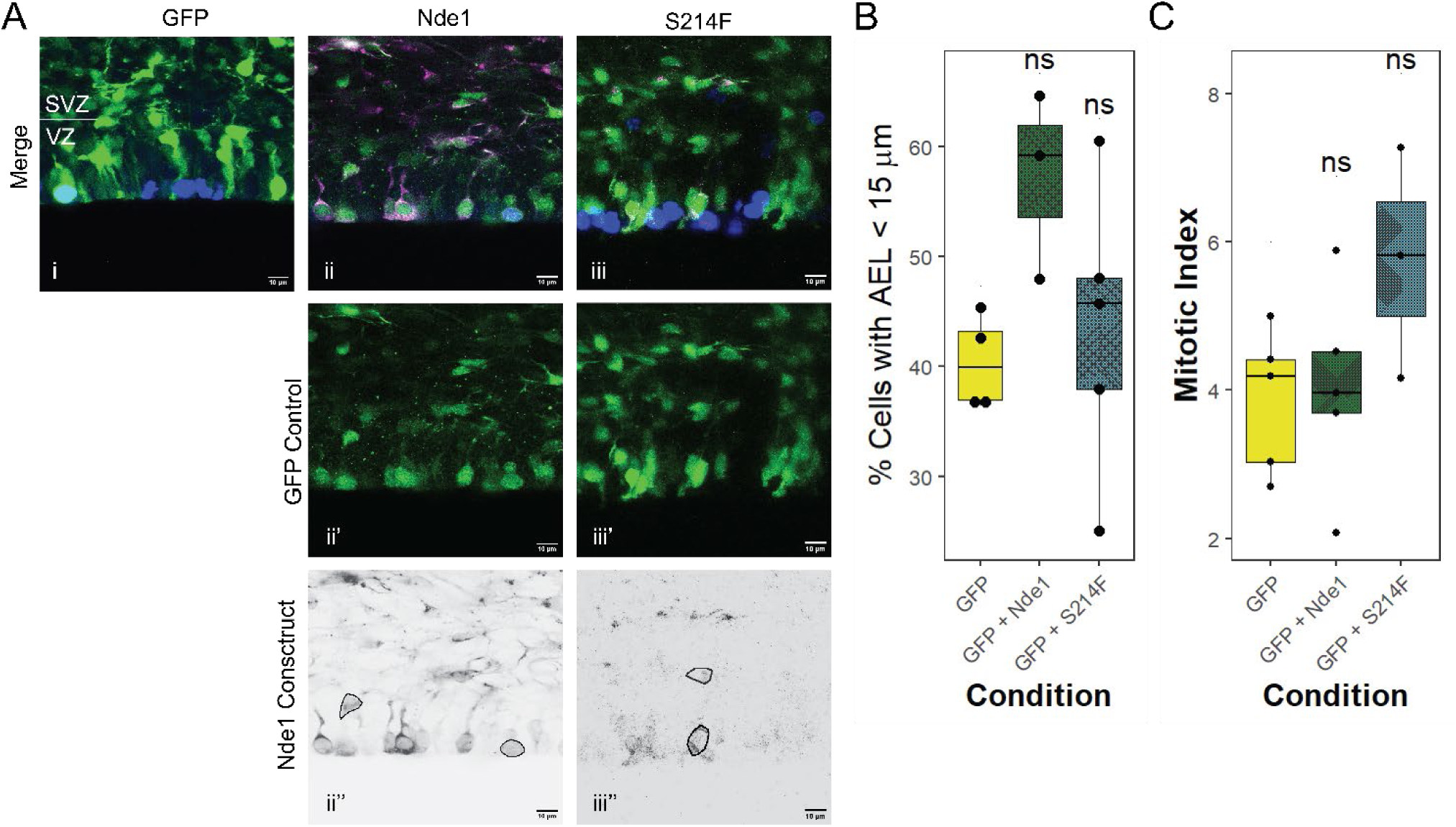
Effects of Schizophrenia associated mutation on RGPs. 6A) Representative images of RGP cells embryonic rat brain subjected to IUE with GFP control plasmid alone (i), Nde1 with GFP control (ii), or S214F with GFP control (iii). In merged images (6Ai-iii), PH3 is shown in blue, GFP in green, and Nde1 constructs in magenta. GFP is shown alone in 6Aii’-iii’. Nde1 or S214F is shown in black in 6Aii’’-iii’’, with example cells outlined in black. Scale bars for all images are 10µm. 6B) Comparison of the percent of cells with an apical endfoot length (AEL) less than 15 µm of the ventricular surface in RGPs expressing GFP alone, and with Nde1 and S214F 6C) Mitotic indices of RGPs expressing GFP alone, and with Nde1 and S214F. All statistical comparisons are made to control data. One-way ANOVA and post hoc Dunnet test were used in 6B-C), ns = p > 0.05. n = mean of values across at least 3 different embryos from at least 2 different operations per condition. All experiments done with E16 operations and E20 analysis in developing rat brains. 6B-C) data plotted as interquartile range with 5-95% whisker range.

Given that Cdk5 is preferentially expressed in post-mitotic neurons,^27^ we next investigated whether the S214F mutation might selectively impair post-mitotic neuronal migration. To test this possibility, we performed *in utero* electroporation with a cDNA encoding Nde1-S214F in a construct containing an IRES sequence and RFP. This was done so the RFP signal would appear throughout the cell, in order to visualize the entire cell morphology of the transfected cells. In addition to a lack of effect on INM, The S214F mutation had no effect on neuronal migration to the cortical plate at E20 (Figure 7A-B). Impaired neuronal migration at E20 might be expected from inhibition of Cdk5 phosphorylation if Nde1 recruitment to the neuronal NE is altered, preventing the cell body from translocating. Importantly, this mutation did alter the morphology of the leading process in in Nde1-S214F-expressing neurons, which showed a significant increase in the percent of cells with branching leading processes, as opposed to a single leading process. This result was also observed following Nde1-T215A expression (Figure 6C-D). We also measured the length of the leading processes in these brains (Figure 7E). Overexpression of Nde1 resulted in an increase in the overall leading process length compared to RFP control, and both T215A and S214F constructs showed significantly reduced leading process length compared to WT Nde1 (Figure 7E). As both mutations, T215A and S214F, show a similar phenotype in E20 neurons, the likely explanation is that Cdk5 phosphorylation at T215 is important for the correct morphology of leading processes. The T215A mutation prevents this by altering the phosphorylated residue, while S214F prevents this by blocking Cdk5 phosphorylation of the adjacent T215, likely through steric hinderance.

**Figure 7.**
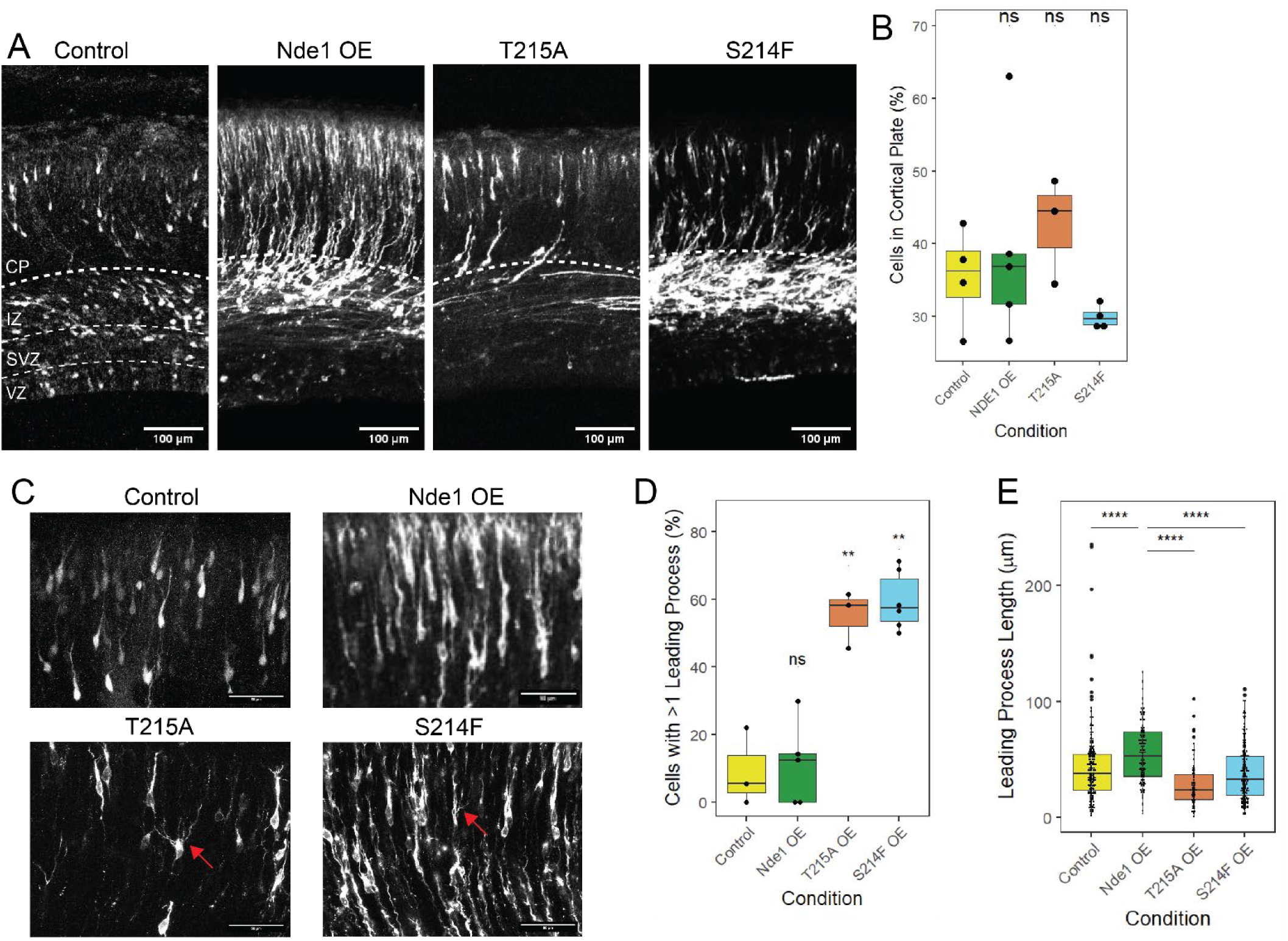
Effects of T214A and S214F Mutations on Neuronal Migration and Morphology. 7A) Representative images of cortices expressing diffuse RFP alone or along with Nde1, T215A, or S214F. In the control image, the four regions of the cortex are indicated by dashed lines. In the other images, only the boundary between the cortical plate (CP) and the rest of the cortex is indicated. Scale bar = 100 µm. 7B) Quantification of the percent of transfected cells that reached the CP in each condition. 7C) Representative images of the neurons in the CP in cells expressing diffuse RFP alone or with Nde1, T215A, or S214F. Red arrows in the T215A and S214F images indicate neurons with >1 leading process. Scale bar = 50 µm 7D) Quantification of the percent of cells with >1 leading process in each condition. 7E) Quantification of the length of leading processes in each condition. Statistical comparisons in 7A and 7D are made to control data and to Nde1 OE in 7E. Unpaired t-test used in 7B and 7D. Kolmogorov-Smirnov test used in 7E. ns = p > 0.05, **p<0.01, ****p<0.0001. n = mean of values across at least 3 different embryos from at least 2 different operations per condition. All experiments done with E16 operations and E20 analysis in developing rat brains. 7B, 7D-E data plotted as interquartile range with 5-95% whisker range.

In order to observe more long-term effects of the S214F mutation, we examined postnatal day 6 (P6) neocortical rat brain slices. These results further supported a neuronal migration defect. Our fixed cell analysis revealed a clear reduction in neuronal number in layers 2/3 of the developing neocortex, as judged by Cux1 staining (Figure 8A-B). WT Nde1 overexpression resulted in a significantly higher percent of cells below layer 2/3 compared to the RFP control, while the T215A and S214F constructs showed a significant reduction in cells below layer 2/3 compared to WT Nde1 (Figure 8B). In contrast, Nde1 KD at P6 results in cells that accumulate primarily within the white matter (Figure 8C) and display proliferative markers that are never seen in the other conditions at P6 (Figure 8D-F).

**Figure 8.**
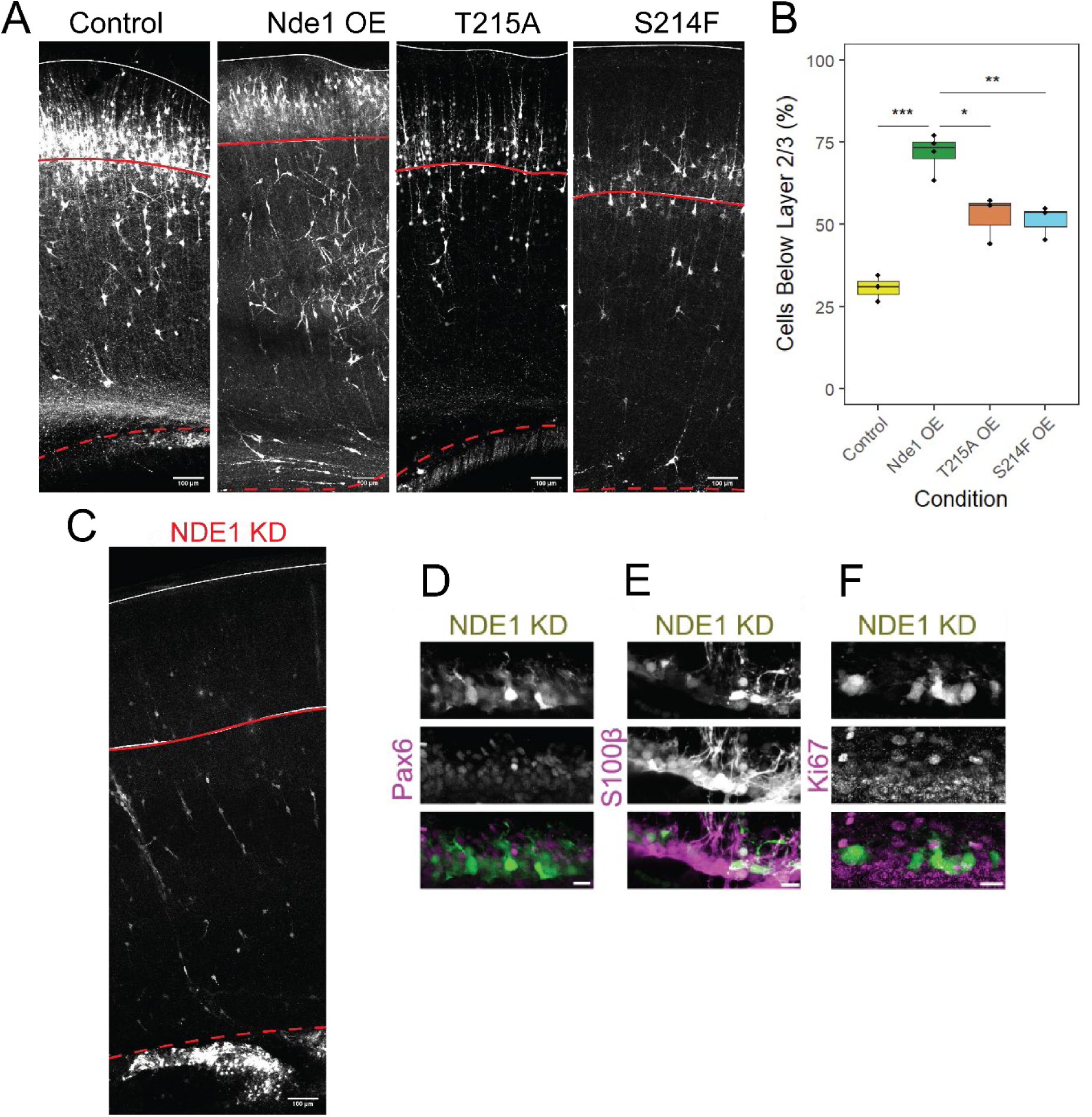
Effects of S214F Mutation on Neuronal Migration in Post-natal Brains. 8A) Representative images of cells expressing diffuse RFP alone or with Nde1 OE, T215A, or S214F in P6 brains. Solid red line indicates the boundary of cortical layer 2/3, which was determined with Cux1 staining. Solid white line indicates the pial surface and red dashed line indicates the boundary of the white matter. Scale bar = 100 µm 8B) Quantification of the percent of cells below the layer 2/3 boundary in each condition. 8C) Representative image of a P6 brain expressing Nde1 KD. Scale bar = 100 µm 8D-F) Representative images of cells in a P6 brain expressing Nde1 KD (green in merged image) as well as proliferative markers Pax 6 (8D), S100β (8E), or Ki67 (8F), shown in magenta in merged images. Scale bar = 10 µm. Unpaired t-test used in 8B. *p<0.05, **p<0.01, ***p<0.001. n = mean of values across at least 3 different brains from at least 2 different operations per condition. All experiments done with E16 operations and P6 analysis. 8B data plotted as interquartile range with 5-95% whisker range.

## Discussion

Considerable progress has been made in understanding the physiological roles of Nde1 and Ndel1 in neocortical development. There is strong evidence for a Nde1 contribution to INM, as well as post-mitotic neuronal migration.^3^ There has been more limited information on the specific roles of Nde1 and Ndel1 phosphoregulation in these processes. We have now tested the contribution of three Nde1 CDK phosphorylation sites in rat brain development. We observed clear and potent effects of specific Nde1 phosphomutants in the control of INM. Our results also provide strong support for new insight into the little-understood mechanism we uncovered for triggering RGP mitotic entry.^3^ Finally, in the course of these studies we obtained clear evidence for a novel post-mitotic role for Nde1 phosphorylation in schizophrenia.

### Differential Roles of Nde1 phosphorylation sites in INM vs. Mitotic entry

Our data serve to validate predicted CDK1 phosphorylation sites at Nde1 residues T215, T243, and T246. We previously evaluated these sites for a role in mitosis in nonneuronal cells and found them to be required for Nde1 binding to the G2 nuclear envelope (NE) as and to prometaphase-to-early anaphase kinetochores in HeLa cells.^15^ *In vitro* analysis performed in that study further revealed that Nde1 phosphorylation enhances its physical interaction with CENP-F, which showed a similar, if not identical, distribution to that of Nde1 in mitotic Hela cells. Here we find a requirement for the same trio of Nde1 sites in RGP mitotic entry in RGP cells. Our current data further indicate that mutations at Nde1 residues T215 and T243 specifically interfere with apical INM in RGPs, presumably due to a failure in recruitment of Nde1 and, in turn, cytoplasmic dynein, to the NE during G2. Given that T215A and T243A each fail to rescue INM in Nde1 RNAi-expressing RGPs, it is likely that both sites must be phosphorylated in order for INM to proceed. We speculate that this might help ensure that mitotic entry cannot occur until RGP cells are sufficiently far along in G2 progression and apical INM.

We find that another Nde1 phosphorylation site, T246, has a novel and distinct function: mutation of this site alone is sufficient to block mitosis without affecting apical INM. It is possible that the T215 and T243 sites must also be phosphorylated for mitotic entry to occur. However, we were unable to test the role of these sites in mitotic entry directly, as their phosphomutant forms arrested cells earlier in G2, during apical INM. Rescue of Nde1 knockdown with the phosphomutant T246A resulted, however, in complete INM, and arrest of RGP nuclei prior to mitotic entry. We do not yet know the specific role of Nde1 in this unexpected mitotic entry control behavior. However, attention to the T246 phosphorylation site may aid further work into mitotic entry requirements of RGPs beyond simple completion of apical INM. A phosphomimetic Nde1 construct is able to rescue INM, but not mitotic entry, in Nde1 KD conditions. This indicates that there may be a specific phosphorylation combination of Nde1 in order to effectively progress through INM and mitosis. These results are modeled in Figure 9. An important note is that in our system, constructs that fail to rescue INM will also fail to rescue mitotic index, as the cells cannot reach the ventricular surface to divide, and we cannot be sure whether these residues have a direct or indirect effect on mitotic entry. Given that the Nde1x3E construct cannot rescue mitotic index in the presence of Nde1 shRNA, it would indicate that either T215 or T243 may need to be dephosphorylated after INM in order to progress through mitosis.

**Figure 9.**
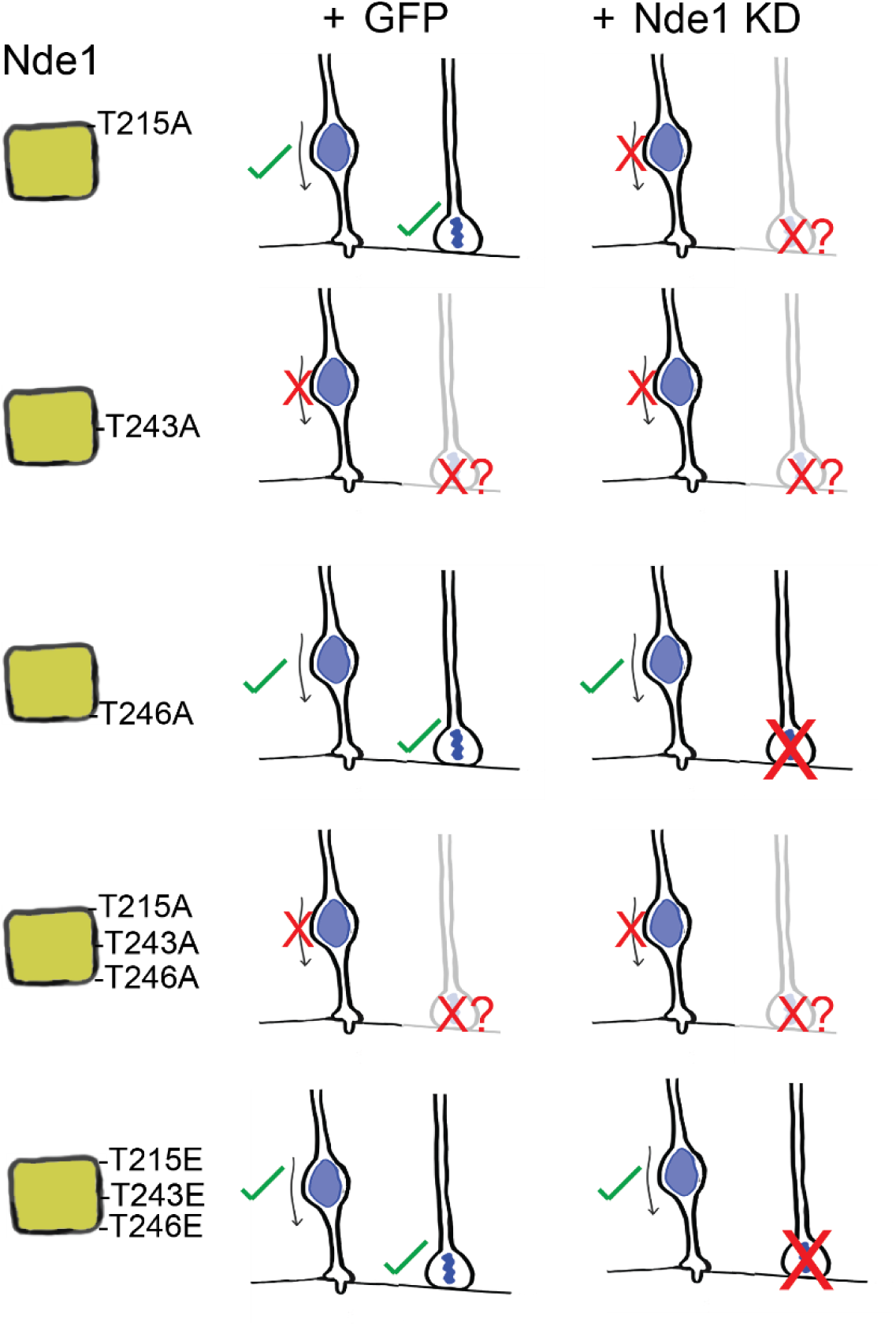
Model of Nde1 phosphorylation sites and their effects on INM and mitotic entry. Out of all constructs tested, only T243A and Nde1x3A had a dominant negative effect on INM. T215A, T243A, and Nde1x3A were unable to rescue INM in combination with Nde1 shRNA. T246A and Nde1x3E were able to rescue INM, but not mitotic index. In the case of constructs that could not rescue INM, mitotic index was affected as RGP nuclei need to reach the ventricular surface in order to divide, but this system does not indicate whether these constructs have a direct effect on mitotic entry.

### Nde1 in schizophrenia

Recent genetic studies have implicated one of the Nde1 phosphorylation sites, Nde1-T215, in schizophrenia. A heterozygous mutation, Nde1 S214F, was found in six unrelated patients with the disease.^22^ This residue is not, itself, phosphorylated by CDK1 or CDK5, but was proposed to interfere sterically with the ability of CDK1 or CDK5 to phosphorylate the adjacent T215 residue. We tested this interesting hypothesis directly, and found it, to be valid: CDK5 phosphorylation of Nde1 was, indeed, inhibited by the S214F mutation, mostly likely through steric hinderance of CDK5 phosphorylation at Nde1-T215. Interestingly, CDK1 phosphorylation was largely unaffected. This remarkable effect seems to be of clear relevance in resolving the molecular basis for the S214F phenotype. In addition to their implications for this particular schizophrenia mechanism, these results indicate how critical phosphoregulation of Nde1 is, and at multiple stages of brain development.

Another important feature of S214F is the nature of the brain developmental phenotype we observe. Unlike the three NDE1 phosphorylation sites we examined, S214F appears to affect post-mitotic aspects of neocortical development in particular. This is consistent with Cdk5 expression in post-mitotic neurons, but not RGPs. We further observed abnormal lamination of Nde1-S214F-expressing cells in the developing neocortex, an unusual, if not unique, anatomic effect of the S214F mutation. This abnormality is similar to the delayed cortical lamination observed in models of Fragile-X syndrome.^28^ Improper neocortical layering could contribute to synaptic abnormalities, which might predispose patients to neuropsychiatric disease. Further investigation of the long-term effects of abnormal neocortical lamination on synaptic development and adult neocortical function will be required to resolve the full consequences of the S214F mutation and its developmental consequences.

## Materials and Methods

### In utero electroporation

Plasmids were transfected by intraventricular injection into the left ventricle of rats at embryonic day 16 (E16) and electroporating the embryos by using a tweezer electrode on either side of the skull outside the external wall of the uterine horn, with the positive electrode adjacent to the injected ventricle. All animal protocols were approved by the Institutional Animal Care and Use Committee at Columbia University.

Cortical slice preparation and immunostaining. 4 days following the in-utero electroporation procedures, the embryos were dissected from the uterus. Embryonic brains were dissected and fixed in 4% paraformaldehyde in PBS for 24 hours at 4°C. Following fixation, the brains were embedded in 4% agarose in PBS and sliced by a vibratome (Zeiss) in 100mm thick coronal sections. The sections were incubated in blocking solution containing PBS with 0.3% Triton X-100 and 5% normal donkey serum for 1 h. Brains were incubated with primary antibodies overnight in blocking solution at 4°C, then sections were washed 3 times in PBS, and incubated in secondary antibodies in blocking solution for 2 h at room temperature. The sections were mounted on slides using Aqua-Poly/Mount (Polysciences, Inc).

### RNAi and DNA constructs

An shRNA construct was designed to target NDE1 and cloned into a pRetro-U6G vector (Cellogenetics, MD, USA), which co-expressed GFP and the shRNA target sequences. The target sequence for NDE1 is 50-GCGTTTGAATCAAGCCAT TGA-30. Empty vectors of pEGFP-C1 and pmCherry-N1 were used as controls (Clontech). For overexpression of NDE1, mouse cDNA for Nde1 was cloned into a mCherry fluorescent reporter (mCherry-C1-NDE1) and subcloned into a pCAGEN vector driven by CAG promoter (provided by Connie Cepko (Addgene plasmid #11160)) using XhoI and NotI restriction sites. Five silent point mutations were made using KOD Hot Start (Millipore) to introduce shRNA resistance in the construct.

### Antibodies

Antibodies used in this study were mouse monoclonal against phosphohistone H3 (Abcam, ab14955, 1:500 dilution), Pax6 (Covance, PRB-278p, 1:300 dilution) S100ß (Millipore, 04-1054, 1:300 dilution), and Ki67 (Millipore, MAB4190, 1:250 dilution) and Cux1 ??. Antibodies were used together with DAPI (4’,6-diamidino-2-phenylindole, Thermo Scientific, 62248, 1:1,000 dilution). NDE1 (Abnova, H00054820-M01, 1:1,000 dilution) and a-tubulin (Sigma; 1:2,000) were used for immunoblotting. To develop in a LICOR system, fluorescent secondary antibodies were acquired from Invitrogen (dilution 1:10,000) and Rockland (dilution 1:10,000) to use for western blotting.

### Imaging and statistical analysis

All images were collected with an IX80 laser scanning confocal microscope (Olympus FV100 Spectral Confocal System). Brain sections were imaged using a 60X 1.42 N.A. oil objective or a 10X 0.40 N.A. air objective. All images were analyzed using ImageJ software (NIH, Bethesda, MD, USA). Apical process length and mitotic index measurements were also performed using this software. All statistical analysis was performed using Prism (GraphPad Software, La Jolla, CA, USA) or R. All brain measurements were made from at least three different embryonic rat brains across at least two mothers. Animals from all successful operations were included in the analysis. For all analyses, significance was accepted at the level of P<0.05.

### Protein purification

Mouse Nde1 cDNA was cloned into the pGEX 6.1 vector using BamHI/NotI sites. Residues Threonine 243 and Threonine 246 were mutagenized into Alanine with KOD polymerase (T243A-T246A). Next, residue Serine 214 was mutated to Phenylalanine with KOD polymerase (S214F). GST-Nde1-T243A-T246A and GST-Nde1-S214F-T243A-T246A constructs were expressed in Rosetta BL21 cells with 0.1mM IPTG at 19°C overnight and purified with GSH beads. 200 ml cultures were lysed in 500 mM NaCl, 50 mM Tris-HCl pH 7.5, 1mM DTT, complete EDTA-free protease inhibitors, 1mM lysozyme, and bound to 600 ul slurry of GSH agarose beads (brand) for 2 h. After 3 washes in 10 bed volumes of 300 mM NaCl, 50 mM Tris-HCl pH 7.5, 1mM DTT, the GST-tagged protein was eluted in 250 mM NaCl, 50 mM Tris-HCl pH 7.5, 1mM DTT, 20 mM reduced GSH, and dialyzed overnight against 150 mM NaCl, 50 mM Tris-HCl pH 7.5, 1mM DTT.

### Kinase assay

10 ug of GST-Nde1-T243A-T246A or GST-Nde1-S214F-T243A-T246A were incubated with or without 0.2 ug active Cdk5 or Cdk1 (Millipore), 400 uM ATP, 10x PK buffer for 45 min at 30C. The kinase reaction was stopped by adding Llaemli loading buffer. A volume of the reaction corresponding to ∼1 ug of GST-tagged protein was loaded on SDS-PAGE gel for western blot analysis. Antibodies used: anti phosphor-Thr(Pro) (Cell Signaling, 1:1000), anti Nde1 (generated as in Stehman et al., 2007)^29^, anti Cdk5 (Santa Cruz, 1: 5000).

## Acknowledgments

The authors thank the members of the Vallee lab for helpful discussions.

## References

1. Tsai, J.-W., Lian, W.-N., Kemal, S., Kriegstein, A. & Vallee, R. B. An Unconventional Kinesin and Cytoplasmic Dynein Are Responsible for Interkinetic Nuclear Migration in Neural Stem Cells. Nat. Neurosci. 13, 1463–1471 (2010).

2. Baffet, A. D., Hu, D. J. & Vallee, R. B. Cdk1 Activates Pre-mitotic Nuclear Envelope Dynein Recruitment and Apical Nuclear Migration in Neural Stem Cells. Dev. Cell 33, 703–716 (2015).

3. Doobin, D. J., Kemal, S., Dantas, T. J. & Vallee, R. B. Severe NDE1-mediated microcephaly results from neural progenitor cell cycle arrests at multiple specific stages. Nat. Commun. 7, 12551 (2016).

4. Reiner, O. et al. Isolation of a Miller-Dieker lissencephaly gene containing G protein beta-subunit-like repeats. Nature 364, 717–721 (1993).

5. Hirotsune, S. et al. Graded reduction of Pafah1b1 (Lis1) activity results in neuronal migration defects and early embryonic lethality. Nat. Genet. 19, 333–339 (1998).

6. Feng, Y. et al. LIS1 Regulates CNS Lamination by Interacting with mNudE, a Central Component of the Centrosome. Neuron 28, 665–679 (2000).

7. Pawlisz, A. S. et al. Lis1–Nde1-dependent neuronal fate control determines cerebral cortical size and lamination. Hum. Mol. Genet. 17, 2441–2455 (2008).

8. Alkuraya, F. S. et al. Human Mutations in NDE1 Cause Extreme Microcephaly with Lissencephaly. Am. J. Hum. Genet. 88, 536–547 (2011).

9. Bakircioglu, M. et al. The essential role of centrosomal NDE1 in human cerebral cortex neurogenesis. Am. J. Hum. Genet. 88, 523–535 (2011).

10. Kriegstein, A. & Alvarez-Buylla, A. The Glial Nature of Embryonic and Adult Neural Stem Cells. Annu. Rev. Neurosci. 32, 149–184 (2009).

11. Hu, D. J.-K. et al. 29_Dynein Recruitment to Nuclear Pores Activates Apical Nuclear Migration and Mitotic Entry in Brain Progenitor Cells. Cell 154, 1300–1313 (2013).

12. Strzyz, P. J. et al. Interkinetic nuclear migration is centrosome independent and ensures apical cell division to maintain tissue integrity. Dev. Cell 32, 203–219 (2015).

13. Splinter, D. et al. Bicaudal D2, Dynein, and Kinesin-1 Associate with Nuclear Pore Complexes and Regulate Centrosome and Nuclear Positioning during Mitotic Entry. PLOS Biol. 8, e1000350 (2010).

14. Splinter, D. et al. BICD2, dynactin, and LIS1 cooperate in regulating dynein recruitment to cellular structures. Mol. Biol. Cell 23, 4226–4241 (2012).

15. Wynne, C. L. & Vallee, R. B. Cdk1 phosphorylation of the dynein adapter Nde1 controls cargo binding from G2 to anaphase. J Cell Biol jcb.201707081 (2018) doi:10.1083/jcb.201707081.

16. Bolhy, S. et al. A Nup133-dependent NPC-anchored network tethers centrosomes to the nuclear envelope in prophase. J. Cell Biol. 192, 855–871 (2011).

17. Wynshaw-Boris, A., Pramparo, T., Youn, Y. H. & Hirotsune, S. Lissencephaly: Mechanistic insights from animal models and potential therapeutic strategies. Semin. Cell Dev. Biol. 21, 823–830 (2010).

18. Hebbar, S. et al. Lis1 and Ndel1 influence the timing of nuclear envelope breakdown in neural stem cells. J. Cell Biol. 182, 1063–1071 (2008).

19. Klinman, E., Tokito, M. & Holzbaur, E. L. F. CDK5-dependent activation of dynein in the axon initial segment regulates polarized cargo transport in neurons. Traffic Cph. Den. 18, 808–824 (2017).

20. Pandey, J. P. & Smith, D. S. A Cdk5-dependent switch regulates Lis1/Ndel1/dynein-driven organelle transport in adult axons. J. Neurosci. Off. J. Soc. Neurosci. 31, 17207–17219 (2011).

21. Blackwood, D. H. R. et al. Schizophrenia and Affective Disorders—Cosegregation with a Translocation at Chromosome 1q42 That Directly Disrupts Brain-Expressed Genes: Clinical and P300 Findings in a Family. Am. J. Hum. Genet. 69, 428–433 (2001).

22. Kimura, H., et al. Identification of Rare, Single-Nucleotide Mutations in NDE1 and Their Contributions to Schizophrenia Susceptibility. Schizophr. Bull. 41, 744–753 (2015).

23. Johnstone, M. et al. Copy Number Variations in DISC1 and DISC1-Interacting Partners in Major Mental Illness. Mol. Neuropsychiatry 1, 175–190 (2015).

24. St Clair, D., et al. Association within a family of a balanced autosomal translocation with major mental illness. The Lancet 336, 13–16 (1990).

25. Millar, J. K. et al. Disruption of two novel genes by a translocation co-segregating with schizophrenia. Hum. Mol. Genet. 9, 1415–1423 (2000).

26. Hirohashi, Y. et al. Centrosomal proteins Nde1 and Su48 form a complex regulated by phosphorylation. Oncogene 25, 6048–6055 (2006).

27. Smith, D. S., Greer, P. L. & Tsai, L. H. Cdk5 on the brain. Cell Growth Differ. Mol. Biol. J. Am. Assoc. Cancer Res. 12, 277–283 (2001).

28. Lee, F. H. F., Lai, T. K. Y., Su, P. & Liu, F. Altered cortical Cytoarchitecture in the Fmr1 knockout mouse. Mol. Brain 12, 56 (2019).

29. Stehman, S. A., Chen, Y., McKenney, R. J. & Vallee, R. B. NudE and NudEL are required for mitotic progression and are involved in dynein recruitment to kinetochores. J. Cell Biol. 178, 583–594 (2007).

